# Controlling homology-directed repair outcomes in human stem cells with dCas9

**DOI:** 10.1101/2021.12.16.472942

**Authors:** William C. Skarnes, Gang Ning, Sofia Giansiracusa, Alexander S. Cruz, Cornelis Blauwendraat, Brandon Saavedra, Kevin Holden, Mark R. Cookson, Michael E. Ward, Justin A. McDonough

## Abstract

Modeling human disease in human stem cells requires precise, scarless editing of single nucleotide variants (SNV) on one or both chromosomes. Here we describe improved conditions for Cas9 RNP editing of SNVs that yield high rates of biallelic homology-directed repair. To recover both heterozygous and homozygous SNV clones, catalytically inactive ‘dCas9’was added to moderate high activity Cas9 RNPs. dCas9 can also block re-cutting and damage to SNV alleles engineered with non-overlapping guide RNAs.

## Main

Editing one or both chromosomes is required to model dominant or recessive disease in human stem cells, respectively. Precise modification of the genome by homology-directed repair can be achieved in human cells with programmable nucleases^1,2^. Diseases are often caused by a single nucleotide variant (SNV) and thus, faithful modeling of pathogenic alleles requires the introduction of a single base change in one or both copies of the target sequence. Here we describe a high-throughput workflow that can be reliably tuned to engineer dominant and recessive disease alleles in human induced pluripotent stem (iPS) cells.

Delivery of Cas9 ribonucleoprotein (RNP), as opposed to plasmids expressing Cas9 and guide RNA, has several advantages for genome editing of cultured cells^3^. Cas9 RNP is immediately active when the donor template is at maximal levels and is degraded within 24 hours, reducing potential off-target damage by avoiding prolonged expression of an active nuclease. Moreover, the amount of Cas9 RNP and repair template taken up by cells can be controlled. Using a BFP reporter assay, we previously demonstrated that Cas9 RNP editing of a single nucleotide variant (H67Y, C to T converting BFP to GFP) is greatly improved by the additive effects of cold shock, a small molecule enhancer of homology-directed repair (HDR), and end-protected donor oligonucleotides^4^. In this study, we employed these conditions to edit Alzheimer’s Disease and Related Dementia (ADRD) variants^5^ in the KOLF2.1JJ reference iPS cell line^6^.

All reagents for editing are available from commercial vendors, including high-fidelity recombinant Cas9 protein, chemically modified single guide RNAs and end-modified donor oligos. Guide RNAs that overlap the SNV were identified and those with the lowest predicted off-target score^7^ were chosen. We avoided altering the obligate ‘GG’ of the PAM sequence to facilitate future reversion experiments using the same PAM. If more than one suitable guide RNA was available, then two guides were tested *in vitro* and the most active guide selected for nucleofection (**Fig. S1**). In cases where *in vitro* activity was similar, we chose guides that direct DNA cleavage nearest to the SNV.

Small volume, parallel nucleofections in a 16-well cuvette were piloted for the first three ADRD models. Genotyping of single cell-derived clones by Sanger sequencing showed efficient HDR; however, the proportion of SNV/WT and SNV/SNV clones was highly variable (**Fig 1a**). Equal numbers of heterozygous SNV/WT and homozygous SNV/SNV clones were recovered for FUS_R495*, whereas SNV/WT clones were preferentially recovered for PFN1_C71G. Remarkably, nearly all of the PFN1_M114T clones were homozygous for the variant and no heterozygous clones were recovered. Lowering the amount of Cas9 RNP reduced the number of PFN1_M114T homozygous clones and overall editing efficiency generally, but failed to produce any SNV/WT clones (data not shown). These early results demonstrate guide-dependent variability in the outcome of CRISPR editing experiments and highlight the challenge of engineering heterozygous SNV/WT clones with highly active Cas9 RNPs.

**Figure 1.**
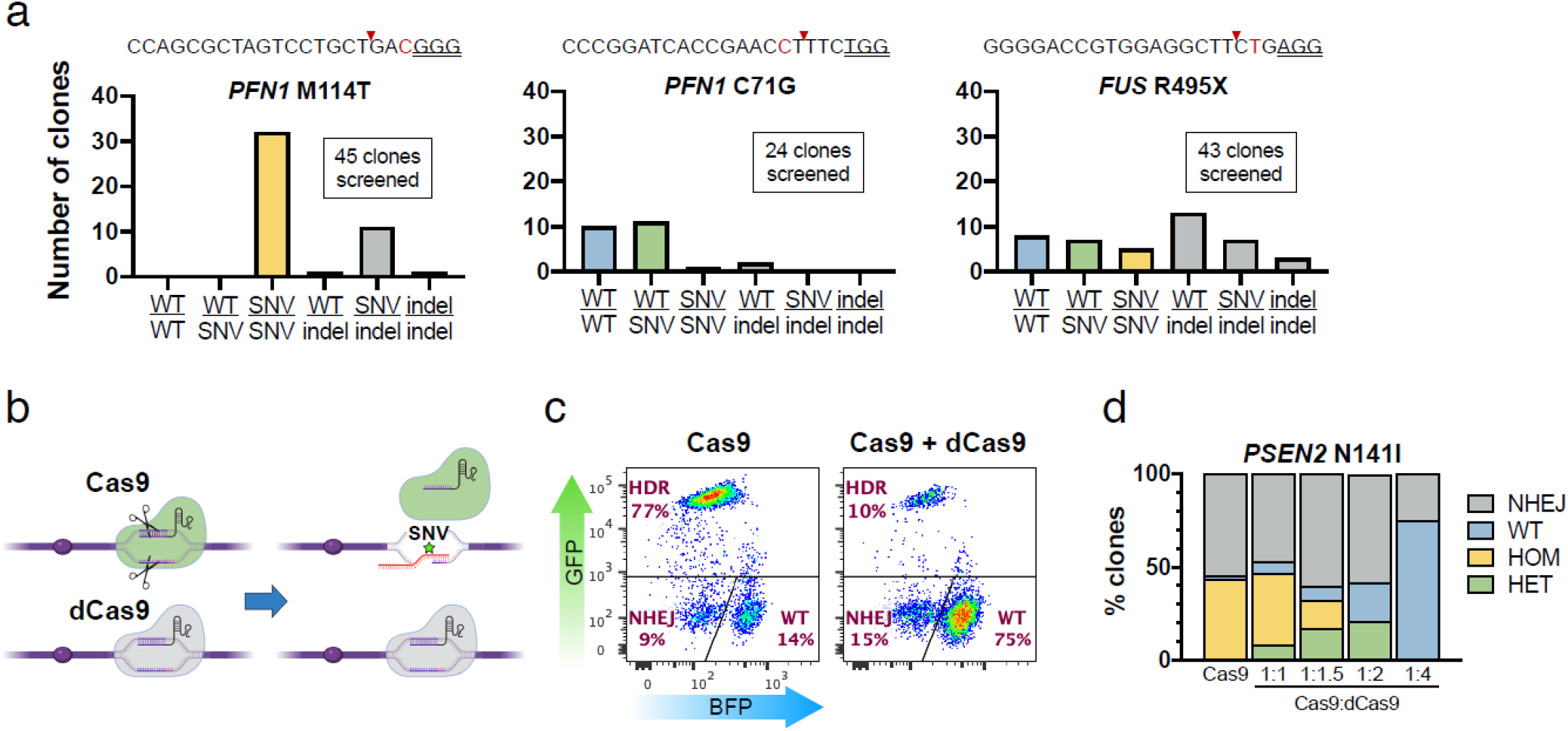
High biallelic editing of SNV alleles using Cas9 RNP and modulation of zygosity with the addition of dCas9. (**a**) Genotyping results for Cas9 RNP editing of three ADRD mutations. Guide RNA sequences overlap the variant (red) and are positioned near the cut site (red triangle), but not within the PAM sequence (underlined). (**b**) Hypothetical modulation of Cas9 RNP activity with dCas9. (**c**) BFP->GFP assay^4^ showing dose-dependent suppression of HDR in the presence of dCas9. (**d**) PSEN2 N141I editing outcomes in the presence of increasing amounts of dCas9.

Nucleofection conditions were further optimized in additional BFP reporter experiments. Maximal HDR efficiency (90% conversion of BFP to GFP) was observed using half the amount of RNP (0.5X), a second generation HDR enhancer and extending the period of cold shock to 3 days (**Fig. S2a**). Conveniently, pre-assembly of Cas9 RNP *in vitro* is not required to achieve optimal HDR (**Fig. S2b**), making it easier to assemble small volume nucleofections. For the next set of 36 ADRD targets, very high editing efficiencies were achieved with this protocol for 90% of the target sequences (**Table 1**).

**Table 1.**
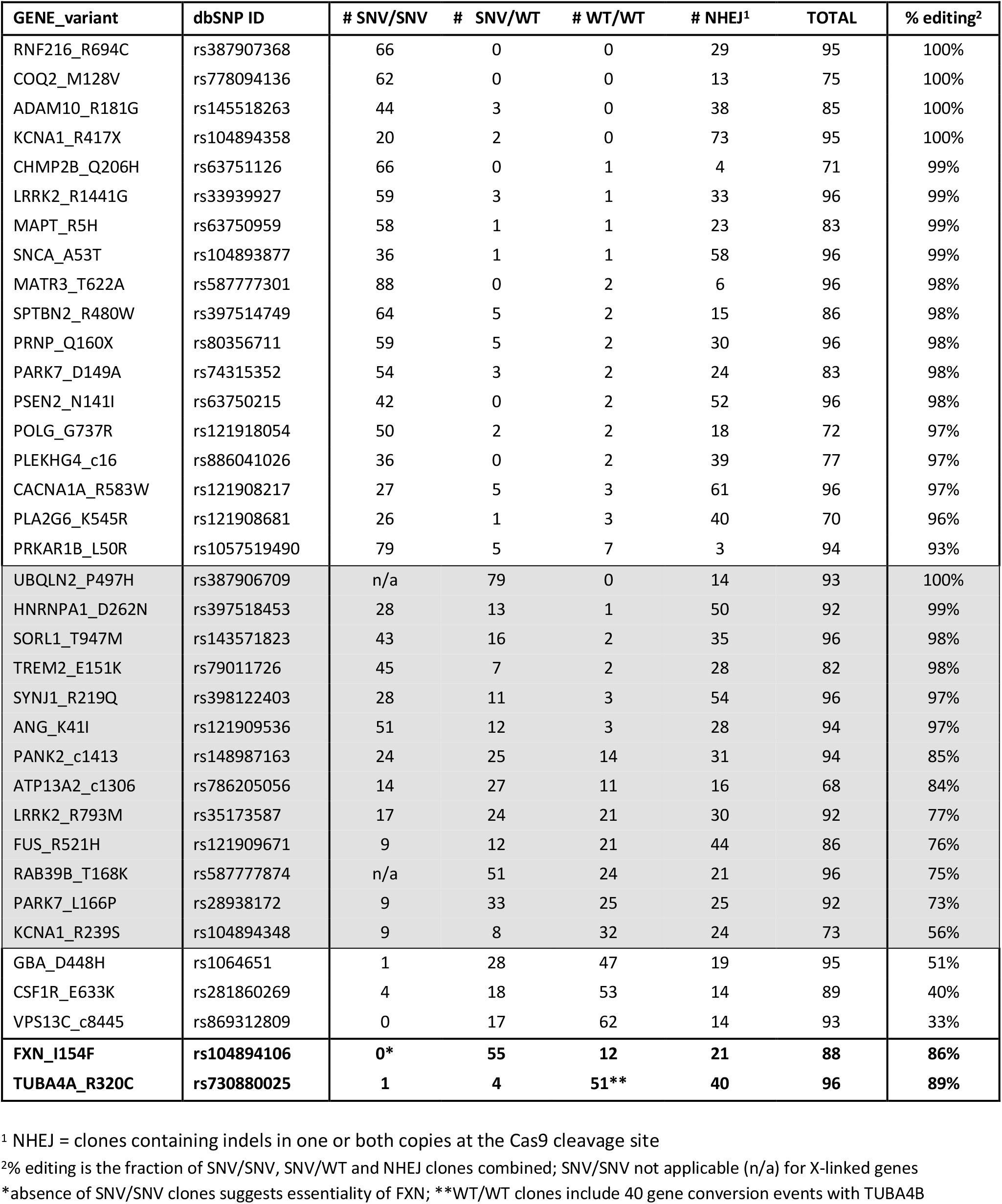
Summary of SNV editing experiments with Cas9 RNP.

| GENE_variant | dbSNP ID | # SNV/SNV | # SNV/WT | # WT/WT | # NHEJ<sup>1</sup> | TOTAL | % editing<sup>2</sup> |
| --- | --- | --- | --- | --- | --- | --- | --- |
| RNF216_R694C | rs387907368 | 66 | 0 | 0 | 29 | 95 | 100% |
| COQ2_M128V | rs778094136 | 62 | 0 | 0 | 13 | 75 | 100% |
| ADAM10_R181G | rs145518263 | 44 | 3 | 0 | 38 | 85 | 100% |
| KCNA1_R417X | rs104894358 | 20 | 2 | 0 | 73 | 95 | 100% |
| CHMP2B_Q206H | rs63751126 | 66 | 0 | 1 | 4 | 71 | 99% |
| LRRK2_R1441G | rs33939927 | 59 | 3 | 1 | 33 | 96 | 99% |
| MAPT_R5H | rs63750959 | 58 | 1 | 1 | 23 | 83 | 99% |
| SNCA_A53T | rs104893877 | 36 | 1 | 1 | 58 | 96 | 99% |
| MATR3_T622A | rs587777301 | 88 | 0 | 2 | 6 | 96 | 98% |
| SPTBN2_R480W | rs397514749 | 64 | 5 | 2 | 15 | 86 | 98% |
| PRNP_Q160X | rs80356711 | 59 | 5 | 2 | 30 | 96 | 98% |
| PARK7_D149A | rs74315352 | 54 | 3 | 2 | 24 | 83 | 98% |
| PSEN2_N141I | rs63750215 | 42 | 0 | 2 | 52 | 96 | 98% |
| POLG_G737R | rs121918054 | 50 | 2 | 2 | 18 | 72 | 97% |
| PLEKHG4_c16 | rs886041026 | 36 | 0 | 2 | 39 | 77 | 97% |
| CACNA1A_R583W | rs121908217 | 27 | 5 | 3 | 61 | 96 | 97% |
| PLA2G6_K545R | rs121908681 | 26 | 1 | 3 | 40 | 70 | 96% |
| PRKAR1B_L50R | rs1057519490 | 79 | 5 | 7 | 3 | 94 | 93% |
| UBQLN2_P497H | rs387906709 | n/a | 79 | 0 | 14 | 93 | 100% |
| HNRNPA1_D262N | rs397518453 | 28 | 13 | 1 | 50 | 92 | 99% |
| SORL1_T947M | rs143571823 | 43 | 16 | 2 | 35 | 96 | 98% |
| TREM2_E151K | rs79011726 | 45 | 7 | 2 | 28 | 82 | 98% |
| SYNJ1_R219Q | rs398122403 | 28 | 11 | 3 | 54 | 96 | 97% |
| ANG_K41I | rs121909536 | 51 | 12 | 3 | 28 | 94 | 97% |
| PANK2_c1413 | rs148987163 | 24 | 25 | 14 | 31 | 94 | 85% |
| ATP13A2_c1306 | rs786205056 | 14 | 27 | 11 | 16 | 68 | 84% |
| LRRK2_R793M | rs35173587 | 17 | 24 | 21 | 30 | 92 | 77% |
| FUS_R521H | rs121909671 | 9 | 12 | 21 | 44 | 86 | 76% |
| RAB39B_T168K | rs587777874 | n/a | 51 | 24 | 21 | 96 | 75% |
| PARK7_L166P | rs28938172 | 9 | 33 | 25 | 25 | 92 | 73% |
| KCNA1_R239S | rs104894348 | 9 | 8 | 32 | 24 | 73 | 56% |
| GBA_D448H | rs1064651 | 1 | 28 | 47 | 19 | 95 | 51% |
| CSF1R_E633K | rs281860269 | 4 | 18 | 53 | 14 | 89 | 40% |
| VPS13C_c8445 | rs869312809 | 0 | 17 | 62 | 14 | 93 | 33% |
| **FXN_I154F** | **rs104894106** | **0\*** | **55** | **12** | **21** | **88** | **86%** |
| **TUBA4A_R320C** | **rs730880025** | **1** | **4** | **51\*\*** | **40** | **96** | **89%** |

<sup>1</sup> NHEJ = clones containing indels in one or both copies at the Cas9 cleavage site
<sup>2</sup>% editing is the fraction of SNV/SNV, SNV/WT and NHEJ clones combined; SNV/SNV not applicable (n/a) for X-linked genes
\*absence of SNV/SNV clones suggests essentiality of FXN; \*\*WT/WT clones include 40 gene conversion events with TUBA4B

Eighteen (50%) of the experiments produced overwhelming numbers of SNV/SNV clones with few or no SNV/WT and WT/WT clones; 13 (36%) targets produced good numbers of both SNV/WT and SNV/SNV clones (or SNV/Y clones for two X-linked genes). Only three targets (8%), excluding FXN and TUBA4A, produced low numbers of SNV/SNV clones and correlated with a lower overall editing efficiency. For such cases, nucleofection of higher amounts of Cas9 RNP may help recover homozygous clones. FXN is a gene required for early post-implantation development in the mouse^8^ and was identified as an essential gene in human stem cells^9^. No homozygous clones were isolated presumably due to deleterious effects of the I154F variant on stem cell viability or self-renewal. Interestingly, gene conversion was the preferred outcome of TUBA4B editing (40 of 96 clones) involving a closely-linked paralog TUBA4B (**Fig. S3**).

We next developed a novel strategy to moderate RNP activity and thus better control the outcome of our editing experiments. We reasoned that the addition of catalytically ‘dead’ Cas9 protein (dCas9) may provide a simple way to block access of active Cas9 to the target sequence (**Fig. 1b**). The effect of dCas9 on editing is sensitive to the amount of guide RNA. Initially, with a molar excess of guide RNA, no effect was observed in the BFP reporter assay using equal amounts of Cas9 and dCas9 RNP (**Fig. S4a**). However, with lower amounts of guide RNA, dCas9 increased the fraction of unedited BFP-positive cells in a dose-dependent manner (**Fig. 1c, Fig. S4b**). Where we observed preferential biallelic editing, we repeated the nucleofections with dCas9. With the addition of a 1:1 – 1:1.5 molar ratio of Cas9:dCas9, the proportion of SNV/WT and SNV/SNV clones improved significantly (**Fig. 1 d, Fig. 2**).

**Fig. 2.**
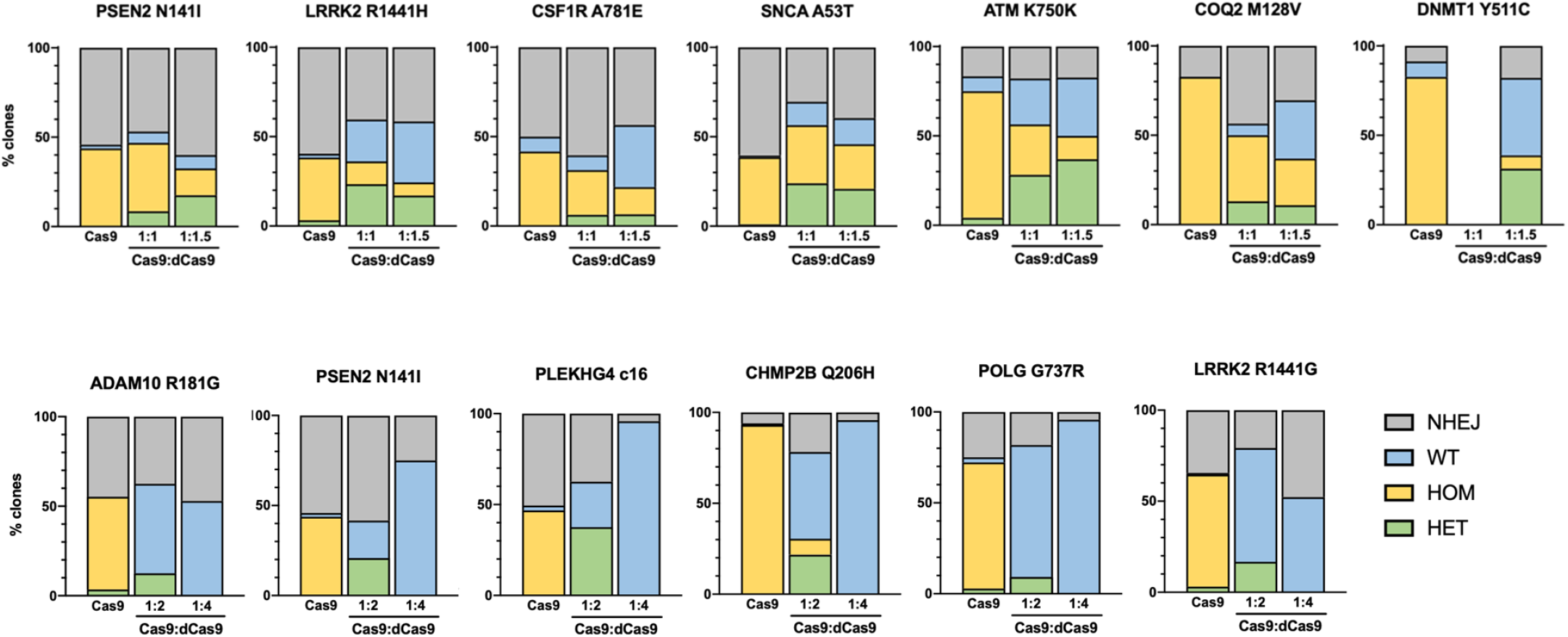
Recovery of SNV/WT clones with dCas9. Editing outcomes for 13 ADRD variants associated with high activity Cas9 RNP showing increasing numbers of SNV/WT and WT/WT clones with increasing amounts of dCas9. At a 1:4 ratio of Cas9:dCas9, HDR is completely suppressed.

Overlapping Cas9 guide RNAs were not available for some SNVs. In such cases, additional silent mutations can be introduced in the repair template to prevent re-cleavage of the modified allele^10^. For modeling human disease alleles, the introduction of additional changes is not ideal, therefore, we explored the application of dCas9 to suppress re-cutting and damage to the modified allele. For two ADRD variants located outside of a suitable guide RNA, nucleofections were performed with and without the addition of dCas9. In the absence of dCas9, all edited alleles were re-cleaved and damaged by error-prone non-homologous end joining (NHEJ; **Fig. S5**), whereas SNV/WT and SNV/SNV clones were readily recovered in the presence of dCas9 (**Table 2**). Further work is required to establish the practical limit of this strategy with respect to the maximum distance of the SNV to the non-overlapping guide.

**Table 2.** Editing of SNVs with non-overlapping guide RNAs.

| GENE_variant | dbSNP ID | Condition | SNV/SNV | SNV/WT | WT/WT | NHEJ* | TOTAL |
| --- | --- | --- | --- | --- | --- | --- | --- |
| SERPINI1_S49P | rs121909051 | Cas9 | 0 | 0 | 0 | 42 | 42 |
|  |  | Cas9+dCas9 | 2 | 9 | 12 | 25 | 48 |
| SETX_L1976R | rs121434379 | Cas9 | 0 | 0 | 4 | 35 | 39 |
|  |  | Cas9+dCas9 | 2 | 8 | 3 | 35 | 48 |

<sup>1</sup> NHEJ = clones containing indels in one or both copies at the Cas9 cleavage site

Our improved editing protocol involves growing cells at 32°C (cold shock) in the presence of a small molecule HDR enhancer. The effect of these conditions on genome stability has not been examined. Damage to off-target sites with 3 or more mismatches in the guide sequence are rare (data not shown); however, ‘on-target’ effects are much more frequent and consequential^11^, involving large indels in one copy of the target sequence or copy-neutral loss-of-heterozygosity (LOH). We analyzed genomic DNA using the NeuroChip DNA microarray^12^ to test for the emergence of aneuploidy, copy number variants or LOH. No detectable copy number changes or aneuploidy was observed in a sample of 96 post-edited clones (8 projects, 6 SNV/WT and 6 SNV/SNV clones per project). Interestingly, two homozygous clones exhibited copy-neutral LOH from the target locus to the distal end of the chromosome (**Fig. S6**).

To more rigorously estimate the rate of ‘on-target’ effects in homozygous SNV/SNV clones we used long-range PCR to amplify a region containing KOLF2.1J-specific heterozygous SNPs on either side of the target site (**Fig. S7**). Clones that exhibit LOH of at least one closely-linked SNP are presumed to have undergone rearrangements (e.g., deletions, insertions, translocations, or inversions) that prevent amplification of one of the two target regions or copy-neutral LOH. The fraction of clones associated with on-target effects varied, ranging from 0% to 13% of clones. Additionally, no chromosome aberrations were found by G-band karyotyping of 32 post-edited clones. Taken together, our QC data show that our protocol for editing human iPS cells has minimal impact on the genome.

In conclusion, we demonstrate the ease by which SNVs can be engineered in human stem cells. Because the PAM sequence is not mutated, each of the alleles described in this study can be reverted to control for any genetic or epigenetic changes that may have arisen in cells during the editing process. We show that dCas9 RNP can be added to moderate highly active guide RNAs and to introduce SNVs outside of the guide sequence. Our fluent, high efficiency workflow is scalable for the production of large numbers of engineered SNVs in one or more reference iPS cell lines and can support organized, systematic phenotyping of human disease models.

## Methods

### Human iPS cell maintenance

KOLF2.1J is an edited iPS subline of KOLF2 (HipSci.org), corrected for a 19 bp deletion in one copy of ARID2^4,6^. Cells were maintained in StemFlex media (Life Technologies) on Synthemax II substrate (Corning) as described^4^ and have undergone extensive characterization^5,6^. After the initial thaw, KOLF2.1J cells were clump-passaged with ReLeSR (Stem Cell Technologies) no more than two times prior to nucleofection.

### Cas9, guide RNA and ssODN

Recombinant HiFi Cas9 v3 (10 ug/ul) and dCas9 (10 ug/ul) were purchased from IDT. Guide RNAs were designed using WGE^7^, modified to display KOLF2.1J variants and Cas12a guide RNAs (wge.jax.org). Guide sequences overlapping the SNV were selected based on two criteria: lowest off-target score and shortest distance of the SNV to the predicted Cas9 cut site. Where available, two sgRNAs per target sequence were tested in an *in vitro* Cas9 digestion assay and the guide with the highest activity was selected for the editing experiment Cas9 sgRNAs, chemically synthesized with 2’-O-methyl and 3’ phosphorothioate end modifications (Synthego CRISPRevolution sgRNA) were resuspended in TE buffer at a concentration of 4ug/ul. End-modified Alt-R™ HDR donor oligos (ssODN, 100-nt) were purchased (IDT) and resuspended in DPBS at a concentration of 200 pmol/ul (see **Supplementary Information** for a complete list of sequences of guide RNAs and oligos).

### *In vitro* Cas9 assay

To test guide RNAs for each target sequence, PCR primer pairs were designed to amplify a 900-1200 bp genomic region containing the CRISPR target site. PCR reactions with 2 ul of purified KOLF2.1J genomic DNA (50 ng/ul) were set up in a 20 ul volume containing 4 ul 5X PrimeSTAR GXL buffer, 0.25 ul of forward and reverse primer (20uM), 1.6 ul of dNTPs (2.5mM of each), 0.5 ul DMSO, 0.4 ul PrimeStar GXL DNA polymerase (1.25 U/ul), and 11 ul HLPC water and 44 cycles (98°C, 10 sec; 60 °C, 15 ses; 68 °C, 2 min) of PCR was performed.

PCR products (150 – 200 ng) were digested for 30 min at 37°C in 20 ul 1X restriction enzyme buffer (NEB buffer v3.1) containing 500 ng Cas9 (Alt-R® S.p. Cas9 Nuclease V3, IDT) and 400 ng sgRNA. (Synthego). The digested products were purified on magnetic beads (AMPure XP) and eluted in 20 ul Tris pH8 according to the manufacturer’s instructions. The relative cutting efficiency of pairs of gRNAs for the same target sequence was visualized and scored by gel electrophoresis (**Fig. S1**).

### High-throughput Cas9 RNP nucleofection

Parallel nucleofections of Cas9 RNP and ssODN in KOLF2.1J cells were carried out in P3 Primary Cell buffer in 16-well cuvettes (Lonza) using an Amaxa 4D-Nucleofector (Lonza). Synthetic sgRNA (Synthego) and end-blocked ssODN (Alt-R HDR Donor Oligos; IDT) were diluted in DPBS to a concentration of 1.6 ug/ul and 40 pmol/ul, respectively, and stored at 4°C for up to several days. Just prior to nucleofection, an 18X master mix of Lonza Primary Cell P3 buffer was prepared fresh by adding 66ul of P3 supplement to 297ul of P3 buffer in a sterile tube. To the P3 Master mix, 3.6 ul Cas9 (10 ug/ul; Alt-R® S.p. HiFi Cas9 Nuclease V3, IDT) was added and mixed thoroughly by pipetting and distributed into sterile 8-well strip tubes (20 ul/tube). 1ul of sgRNA/ssODN was then added to the master mix, such that each nucleofection contains 2ug Cas9, 1.6 ug sgRNA (1:4 Cas9:sgRNA molar ratio) and 40pmol ssODN. For experiments using both Cas9 and dCas9, 2 ug (1:1 Cas9:dCas9), 3ug (1:1.5), 4 ug (1:2) or 8 ug (1:4) of dCas9 (Alt-R® S.p. dCas9 Protein V3, IDT) was added to 20 ul of P3 Master/Cas9 mix containing 2 ug HiFi Cas9 and 0.4 ug sgRNA (1:1 Cas9:sgRNA molar ratio).

Two near-confluent wells of a 6-well plate contain sufficient numbers of cells for one 16-well nucleofection. To prepare a single cell suspension, each well was treated with Accutase and scraped into 2 ml of StemFlex media containing 1X Revitacell (Life Technologies). 1.6×10^5^ cells/well was transferred to two columns of a 96-well V-bottom tissue culture plate and pelleted at 300xg for 5 min at RT. With an 8-channel pipetman, the media was removed and cell pellets were resuspended in 20 ul of the P3/Cas9 RNP/ssODN mix. The cell suspension was transferred to a 16-well cuvette and electroporated using Amaxa program CA-137. Immediately following electroporation, 160 ul of StemFlex media containing 1X Revitacell and 1uM final Alt-R™ HDR Enhancer V2 (IDT) was added to cells and one-third (60ul) was distributed to duplicate wells of a Matrigel-coated 96-well plate. The cells were cultured under ‘cold shock’ conditions in a 32°C/5% CO_2_ incubator for 3 days then returned to 37°C. At 24h post-nucleofection, and every other day thereafter, the media was replaced with StemFlex only. Upon reaching 80% confluency, cells were dissociated to single cells with Accutase and cryopreserved in Knockout Serum Replacement containing 10% DMSO as described previously^4^.

### Single cell cloning

Edited cell pools were thawed in batches on a Matrigel-coated 24-well plate (one pool per well) and grown for several days. Upon near confluency, cells were dissociated to single cells with Accutase (10 min at 37°C), pelleted for 3 min at 300xg, resuspended in 1ml StemFlex with RevitaCell, and counted with a haemocytometer. The single cell suspension was diluted 1:20 in media and 1,500 cells were transferred to a vitronectin-coated 10cm dish containing 10ml StemFlex with 1X RevitaCell. Media (StemFlex only) was replaced one day after plating and every other day thereafter until visible iPS cell colonies formed around day 10. Colonies were manually picked as described previously^4^ and incubated in replicate Matrigel-coated 96-well plates for 4-5 days. Cells from one of the two plates were archived in freezing media in a 96-well Matrix plate^4^. Cells from the duplicate plate were washed with DPBS and frozen without liquid at −80°C for subsequent isolation of genomic DNA.

### Genotyping

Lysis buffer containing Proteinase K was prepared fresh and 100ul per well was added immediately to the frozen 96-well plate of cells as described previously^4^. After a 4h incubation at 60°C, lysates containing genomic DNA were transferred to a 96-well PCR plate while still warm and incubated for 10 min at 95°C in a deep well thermocycler to inactivate Proteinase K. Lysates were diluted 1:10 - 1:20 in water and 2ul was used as a genomic DNA template in a 20 ul PCR reaction, described above.

PCR primer pairs were designed to amplify a 900-1200 bp genomic region containing the CRISPR target site. A third primer was designed for Sanger sequencing of the CRISPR target site. Sanger sequencing of bead-purified PCR products was performed (Quintara) and traces were aligned using SeqMan (DNAStar) or Synthego ICE software (ice.synthego.com).

Analysis of genomic DNA on NeuroChip arrays was performed as described^6^. In brief, genomic DNA isolated from iPS cell lines was genotyped using the NeuroChip array with standard Illumina protocols^12^. To assess genomic integrity of iPS lines we investigated the B-allele frequency and Log R ratio values of each iPS line which were downloaded from Illumina GenomeStudio. Abnormal patterns were observed by visual inspection and were plotted using R (v3.6.1).

## Acknowledgements

We gratefully acknowledge the contribution of Scientific Services at The Jackson Laboratory for expert assistance with the work described in this publication, including Cellular Engineering, Genome Technologies, Flow Cytometry, and Scientific Instrument services. We also thank Glen Beane (The Jackson Laboratory) for modifications to the WGE software; Garret Rettig and Ashley Jacobi (IDT) for HDR enhancer V2; and Haifeng Wang (Tsinghua University) for a stimulating discussion. This work was supported in part by the National Institutes of Health, NIA, Intramural Research Program and by the Jackson Laboratory.

---

**Fig. S1.**
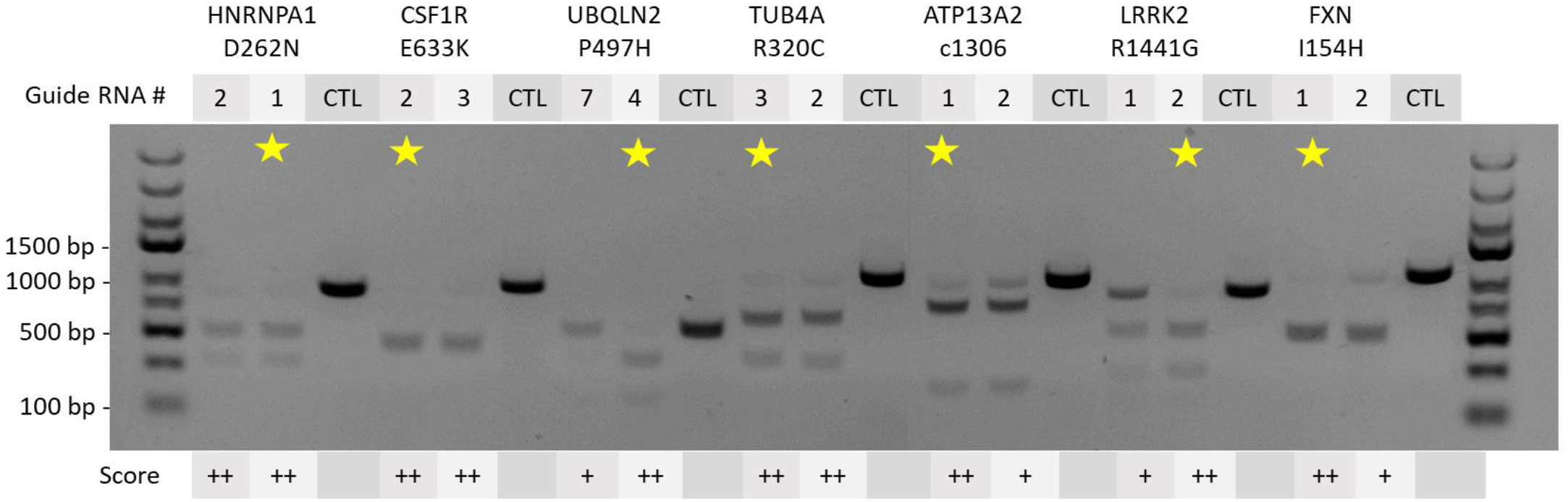
Comparison of the cutting efficiency of pairs of overlapping guide RNAs for each ADRD target sequence. *In vitro* cleavage by Cas9 RNP of PCR-amplified target sequences was visualized by gel electrophoresis. CTL, undigested PCR products. Guide RNAs selected for editing experiments are starred.

**Fig. S2.**
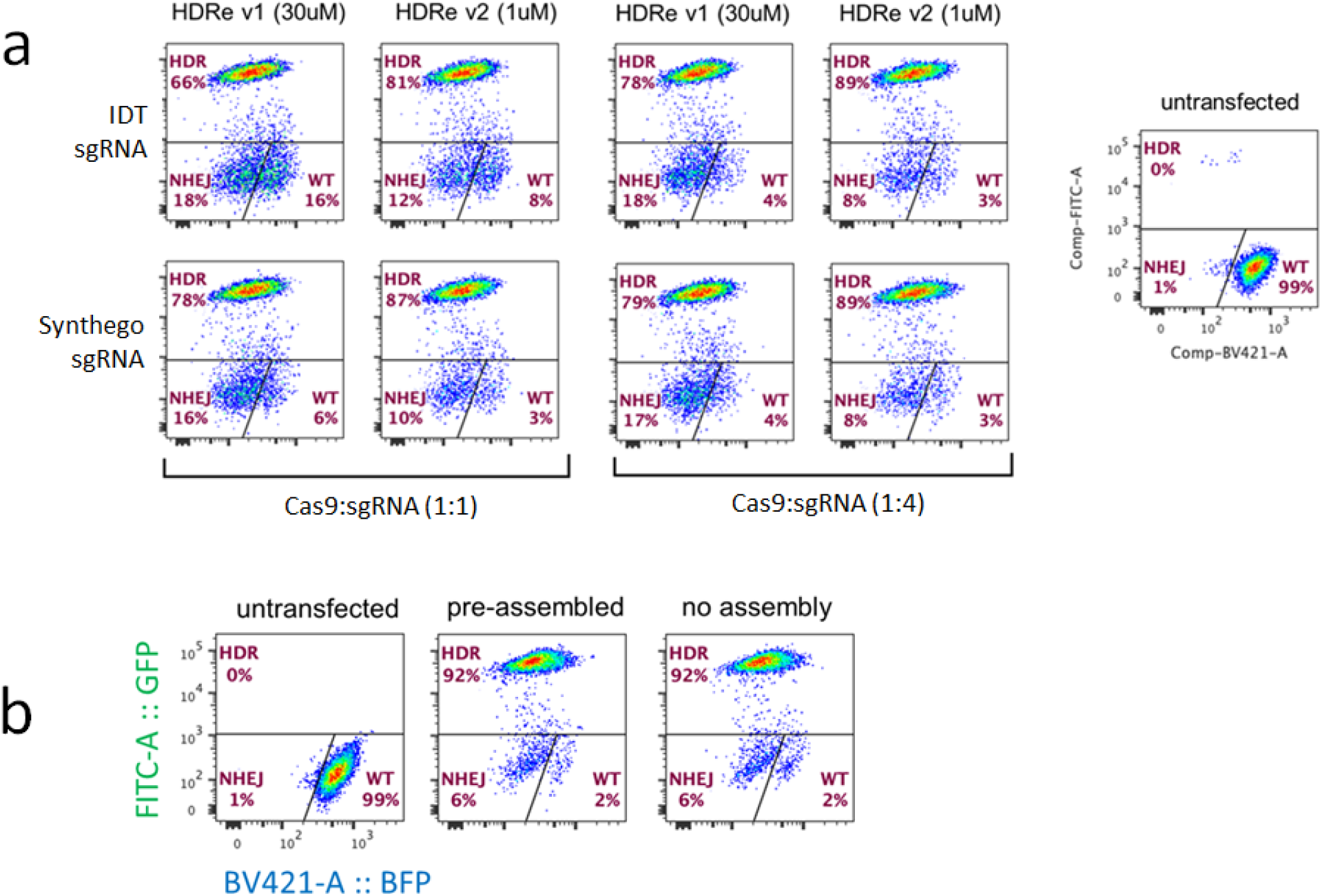
Optimization of HDR. (**a**) BFP reporter assay^4^ comparing various reagents and conditions on HDR efficiency. Synthego guide RNA is more efficient than IDT guide RNA at a 1:1 molar ratio of Cas9:sgRNA. A higher efficiency of HDR was observed in this assay with iDT HDR enhancer V2 compared to HDR enhancer V1. (**b**) Pre-assembly of Cas9 RNP *in vitro* is not required for maximal HDR efficiency.

**Fig. S3.**
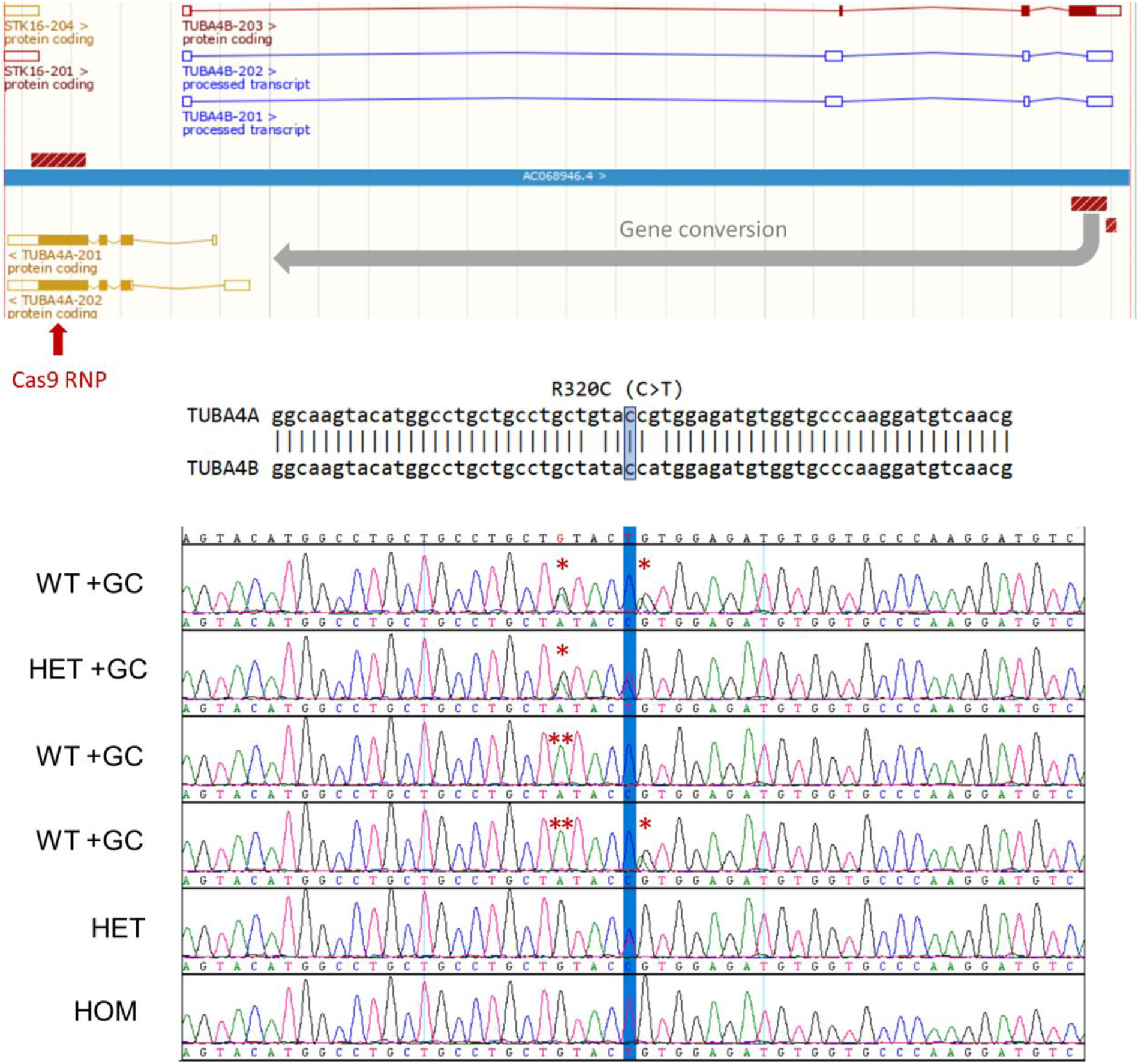
Gene conversion is the preferred outcome for TUBA4B_R320C editing. (Top) Ensembl view of TUBA4A and TUBA4B genes. Hatched red boxes indicate region of high sequence similarity. Cas9 RNP, specific for TUBA4A, induces a double strand break which is repaired by gene conversion in most clones. (Bottom) Sanger sequencing showing the genotypes of edited clones. In 40 of 96 colonies screened, gene conversion (GC) events incorporated TUBA4B-specific sequence on one (*) or both (**) strands.

**Fig. S4.**
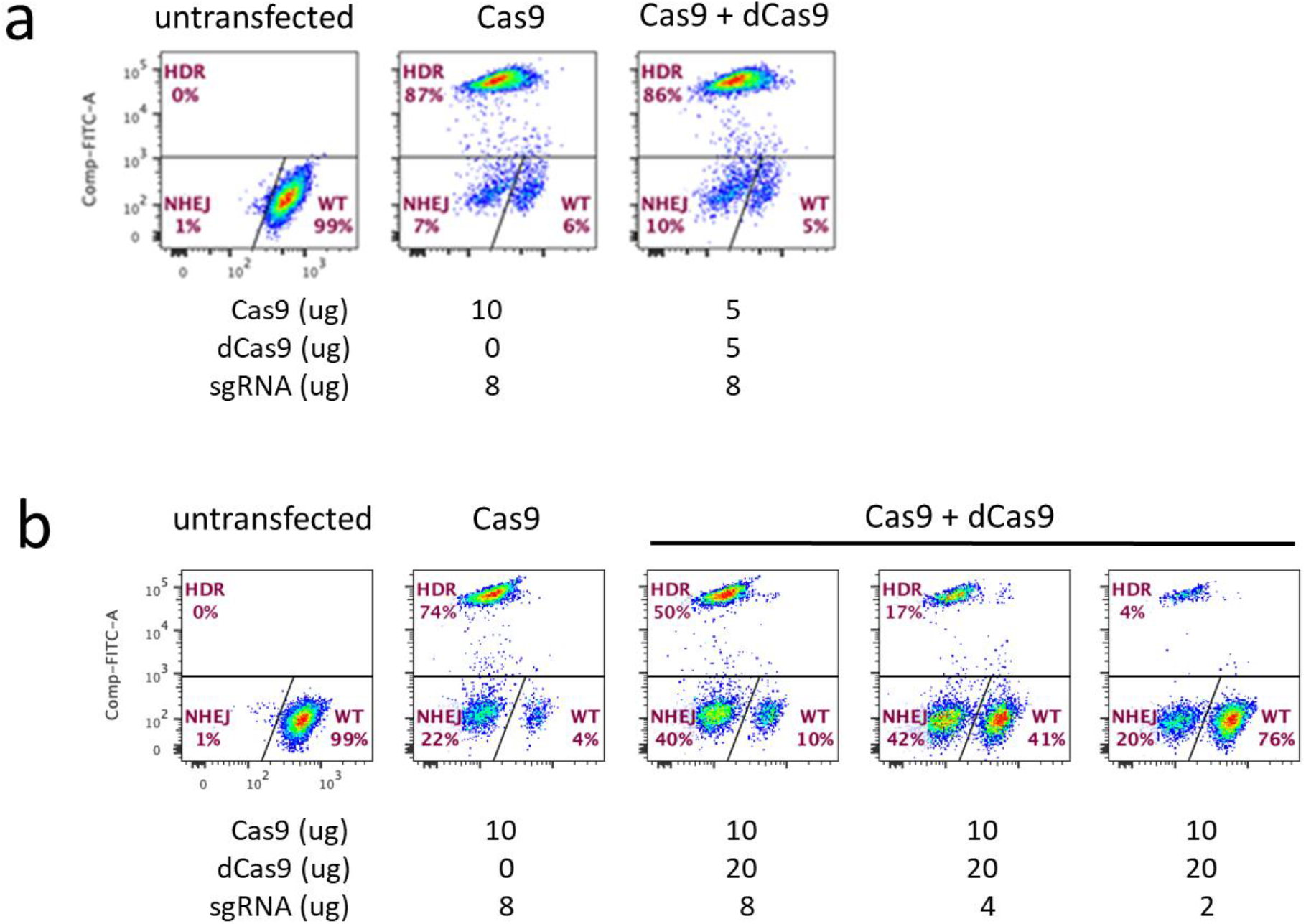
BFP reporter assay^4^ to test the effect of dCas9 on HDR. (**a**) Flow cytometry of nucleofected cells showing no effect on HDR using equimolar amounts Cas9 and dCas9 in the presence of a 4:1 molar excess of guide RNA. (**b**) Dose-dependent suppression of HDR with addition of dCas9 and decreasing amounts of guide RNA.

**Fig. S5.**
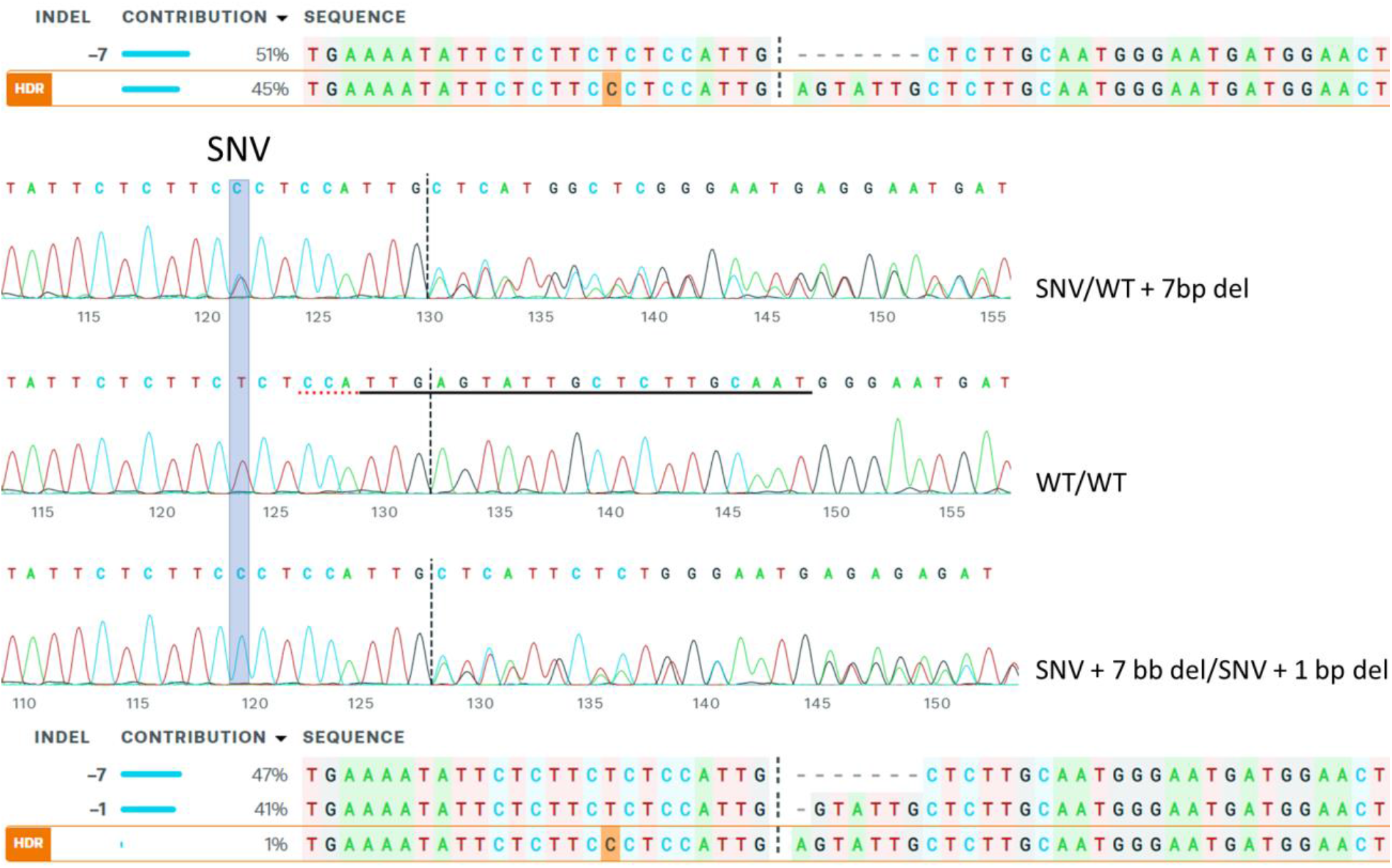
Editing SNV alleles with non-overlapping guide RNAs. Sanger sequencing and ICE analysis shows that all SNV clones also contain indels in one or both copies of the target sequence. The highlighted base change for SERPINI1 S49P variant (T>C) is located outside of the guide sequence (underlined) and PAM sequence (dotted red line), 9 bases from the Cas9 cleavage site (dotted line).

**Fig. S6.**
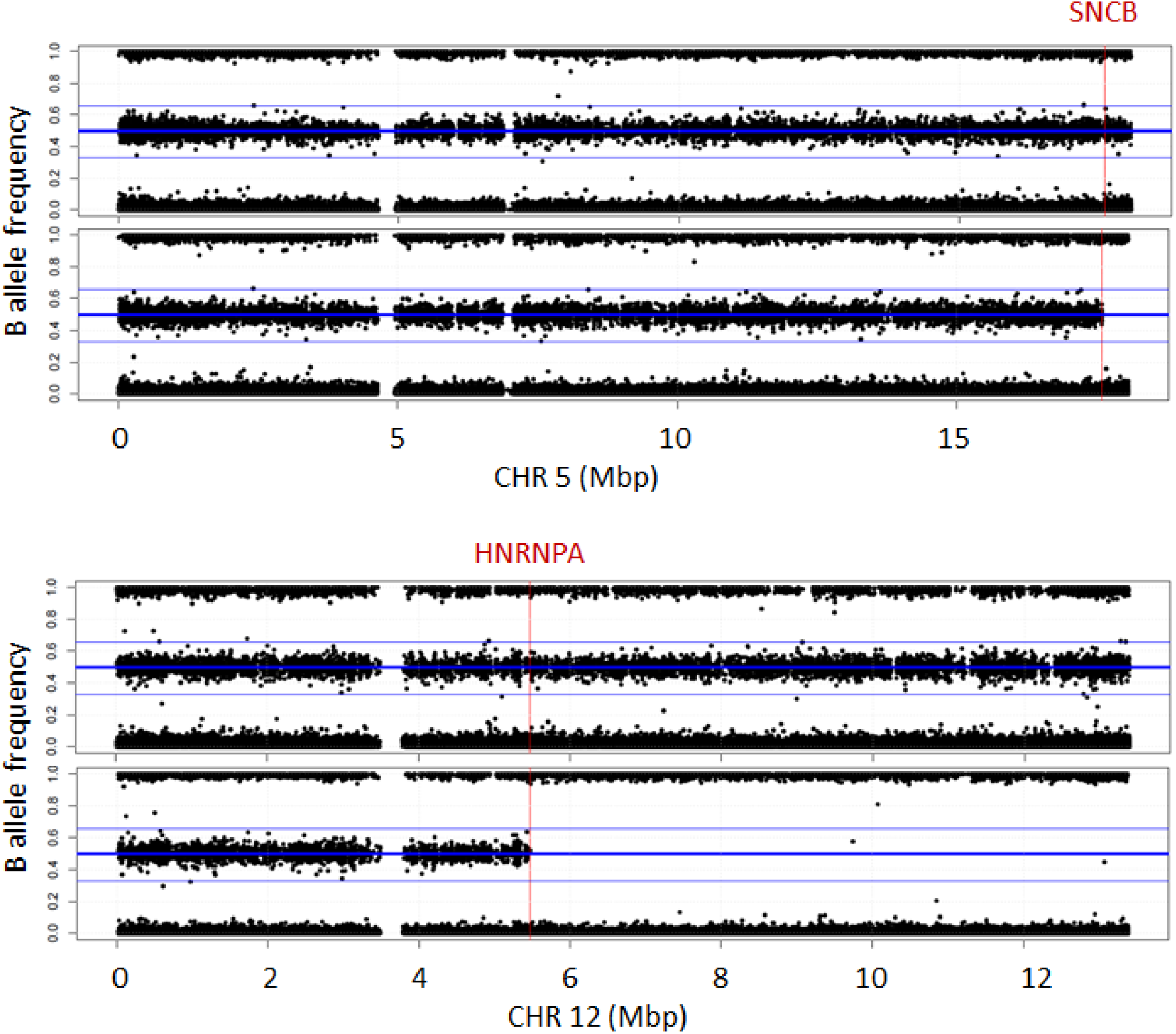
Copy neutral loss-of-heterozygosity detection on NeuroChip arrays, extending from the target locus (red line) to the distal end of the chromosome, detected in a homozygous clone for SNCB P123H (Chr5) and for HNRNPA1 D262N (Chr12).

**Fig. S7.**
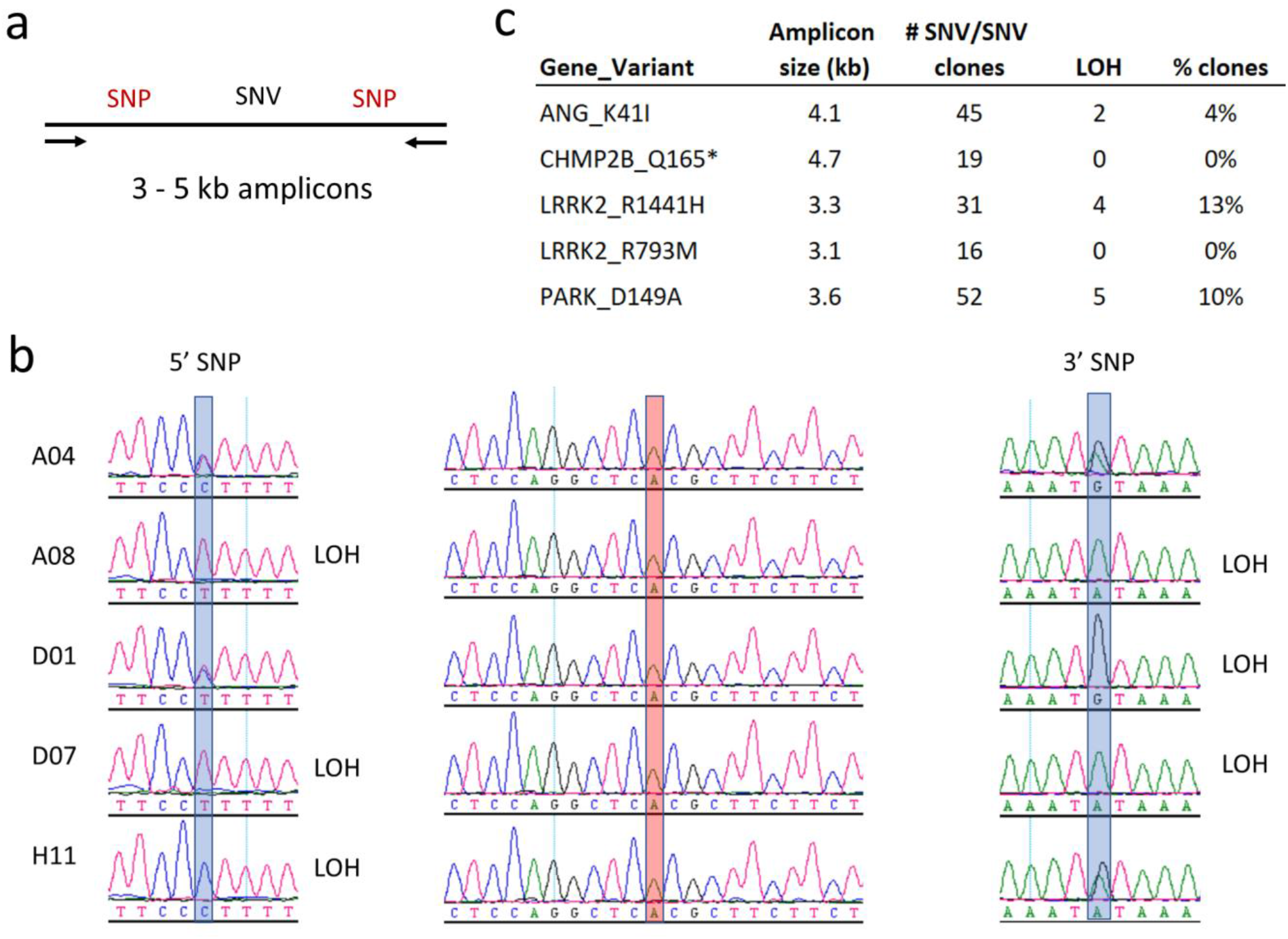
Detection of ‘on-target’ events by long-range PCR and Sanger sequencing. (**a**) PCR amplicons, are designed to include KOLF2.1J heterozygous SNPs closely linked to the SNV. (**b**) Homozygous LRKK2_R141H clones showing LOH of SNPs on one or both sides of SNV (**c**) Summary of SNP genotyping results for five ADRD projects.

